# MutSignatures: An R Package for Extraction and Analysis of Cancer Mutational Signatures

**DOI:** 10.1101/2020.03.15.992826

**Authors:** Damiano Fantini, Vania Vidimar, Yanni Yu, Salvatore Condello, Joshua J. Meeks

## Abstract

Cancer cells accumulate somatic mutations as result of DNA damage and inaccurate repair mechanisms. Different genetic instability processes result in distinct non-random patterns of DNA mutations, also known as mutational signatures. We developed *mutSignatures*, an integrated R-based computational framework aimed at deciphering DNA mutational signatures. Our software provides advanced functions for importing DNA variants, computing mutation types, and extracting mutational signatures via non-negative matrix factorization. We applied *mutSignatures* to analyze somatic mutations found in smoking-related cancer datasets. We characterized mutational signatures that were consistent with those reported before in independent investigations. Our work demonstrates that selected mutational signatures correlated with specific clinical and molecular features across different cancer types, and revealed complementarity of specific mutational patterns that has not previously been identified. In conclusion, we propose *mutSignatures* as a powerful open-source tool for detecting the molecular determinants of cancer and gathering insights into cancer biology and treatment.

## INTRODUCTION

Genetic instability is one of the hallmarks of cancer (1). Neoplastic cells accumulate somatic mutations in their genomes, resulting in aberrant homeostasis, cancer cell survival, and proliferation (2). DNA mutations are the result of the unbalanced interplay between processes generating nucleotide lesions and impaired activity of DNA repair pathways (3). Often, specific mutations can be traced back to the genetic instability process that generated them. For example, 8-oxoguanine (8-oxoG) is the most common and best-characterized base lesion induced by oxidative stress (4,5), a condition associated with cancer (6). During DNA replication, 8-oxoG can pair with adenine, causing G→T transversions (4,7). On the contrary, UV radiation elicits C→T substitutions at dipyrimidine sites, inducing CC→TT (8). Likewise, other molecular processes can be associated with their cognate mutational signatures. The interest in the identification of mutational signatures and the corresponding genetic instability processes is rapidly growing because these signatures are footprints of the molecular aberrations occurring in tumors, may be prognostic of clinical outcomes, and could support personalized anti-cancer treatments in the future (9).

Seminal work from Nik-Zainal et al (10) and Alexandrov et al (11) identified a list of 30 tri-nucleotide mutational signatures found in human cancer (Catalogue of Somatic mutations in Cancer, COSMIC signatures). The analytic pipeline was written in *MATLAB* (Wellcome Trust Sanger Institute, WTSI framework), and relied on non-negative matrix factorization (NMF) (12). NMF has been widely employed to learn the basic components of objects that can be represented as non-negative numeric matrices (13,14), such as mutation counts. Analyses aimed at deciphering mutational signatures were also performed using R-based pipelines and the NMF package (15-17). In addition, R packages dedicated to the identification of tri-nucleotide mutational signatures by NMF and PCA (*somaticSignatures* R package) (18), or using original probabilistic models (*pmsignature* R package) (19) were published. However current R-based approaches for mutational signature analysis carry a series of limitations. First, most analytic pipelines lack built-in functionalities for computing tri-nucleotide mutations, or only support analysis of human mutations. Second, with few exceptions, tri-nucleotide mutations are the only types of DNA variants that were analyzed, even if recent reports suggested that the standard tri-nucleotide-based approaches may be inadequate to capture and resolve clinically- or biologically-relevant patterns. For example, it was recently shown that incorporating additional mutation-flanking nucleotides could be advantageous for better establishing mutational blueprints of smoke-associated cancers (17). Additionally, current approaches are limited by both reproducibility issues emerging when comparing results from different signature extraction pipelines, as well as biases due to differences in total mutation burden across sequenced samples (20). Finally, a fully integrated R-based framework for the analysis of DNA variants and the identification and analysis of mutational signatures is still missing.

These considerations prompted us to develop a software that replicated the *WTSI* framework in the R Statistical Computing environment, and at the same time addressed some of the limitations of the current analytical approaches. Here, we present *mutSignatures*, which is available on CRAN (https://CRAN.R-project.org/package=mutSignatures) and GitHub (https://github.com/dami82/mutSignatures). This framework includes an R-ported version of the software developed by Alexandrov et al (12), accompanied by a wide set of functions for data import, preparation, analysis, and visualization. Notably, our software is compatible with non-human genomes, and was successfully employed to extract for the first time two mutational signatures from a carcinogen-induced mouse model of bladder cancer (21). Moreover, *mutSignatures* provides users with optional tools for inspecting non-standard mutation types, applying sample-wise mutation count normalization, and using a multiplicative update NMF algorithm (22) alternative to the standard Brunet’s algorithm (13). Altogether, *mutSignatures* is a powerful open-source framework for comprehensive analysis of mutational signatures, aimed at gathering insights into cancer biology and treatment.

## MATERIAL AND METHODS

### Data sources

LUAD and BLCA TCGA datasets were described before (23,24). The MAF files storing mutation data from sequencing experiments were downloaded from the Broad Institute Repository at the following URL: http://gdac.broadinstitute.org/runs/analyses__2016_01_28/reports/cancer/. Tri-nucleotide mutation frequencies of 30 COSMIC signatures were downloaded from the Sanger Institute repositories, at the following URL: http://cancer.sanger.ac.uk/cancergenome/assets/signatures_probabilities.txt. The TCGAretriever (https://CRAN.R-project.org/package=TCGAretriever) R package was used to download patient clinical data from cBioPortal (http://www.cbioportal.org).

### Computing Mutation types

*mutSignatures* version 1.3.7 or higher (https://github.com/dami82/mutSignatures) was used. Tri-nucleotide or non-standard mutation types were computed starting from MAF files, and using *mutSignatures* functions that relied on the use of *GenomicRanges* (25) and the *BSgenome* (https://doi.org/doi:10.18129/B9.bioc.BSgenome.Hsapiens.UCSC.hg19) *Bioconductor* packages. Specifically, the full genome sequences for Homo Sapiens, version *hg19* were used for retrieving the nucleotide context surrounding each SNV in the MAF files, and for computing mutation types. Reverse-complement transformations were applied to format all mutations according to the standard style used by COSMIC, which always lists a pyrimidine as the reference base at the mutated position.

### Non-negative Matrix Factorization

The core functions for performing NMF were ported into R from the *MATLAB*-based code of the WTSI (recently renamed to *sigProfiler*) framework (12), which was downloaded from the following URL: https://www.mathworks.com/matlabcentral/fileexchange/38724. NMF was performed using matrix algebra functions that are included in R base. The Brunet’s and the Lin’s NMF algorithms were described before (13,22), and the corresponding MATLAB code (12,22) was ported to R. *De novo* signature extractions by NMF were performed by running at least 500 iterations, and using on-demand Amazon (Seattle, WA, USA) Elastic Cloud 2 (EC2) Linux instances, typically equipped with at least 36 CPU cores and 60 Gb RAM (*m4.10xlarge*, or *c4.8xlarge* EC2 instances).

### Simulations, statistical analyses, and patient prognosis

All statistical tests and data analyses were performed using R. Patient survival analyses were performed using the *survival* R package (https://CRAN.R-project.org/package=survival). For analysis of clinical prognosis in the LUAD dataset, patients were assigned in 2 groups: cases with survival time longer than 36 months were included in the first group (good prognosis, n=111), while deceased patients with survival time shorter than 36 months were included in the second group (poor survival, n=111). Patients with insufficient follow-up time (survival status = alive & survival time less than 36 months; n=196) were excluded from the ‘prognosis’ analysis.

Signature matching was performed using the *matchSignatures()* function from the *mutSignatures* package. This function computed the cosine distance of all pairs of signatures from two *mutationSignatures* objects (dist=0 meant identity; dist∼1 meant maximum dissimilarity). Results were visualized by heatmaps.

For the Monte Carlo simulation, a total of 10,000 simulations were performed. At each iteration, relative signature exposures of 418 genomes were generated, so that each signature had relative exposure distribution whose mean and standard deviation tracked with those observed in the original signature exposures. Spearman correlation was then computed for all pairs of signatures, and the minimum correlation value was returned. Finally, the original correlation values were examined with respect to the distribution of correlation values returned by all simulations. Spearman’s and Kendall’s correlation tests were performed using the *cor.test()* function from the *stats* R package.

## IMPLEMENTATION

### Overview of mutSignatures pipeline

The *mutSignatures* framework is organized in three modules (figure 1). The first module deals with data import and preparation from Variant Call Format (VCF) files or other sources. The second module includes core functions required for *de novo* extraction of mutational signatures by NMF. Alternatively, mutation counts can be deconvoluted against known mutational signatures to determine signature exposures. The third module includes functions for mutational signature matching, downstream analysis, and visualization.

**Figure 1.**
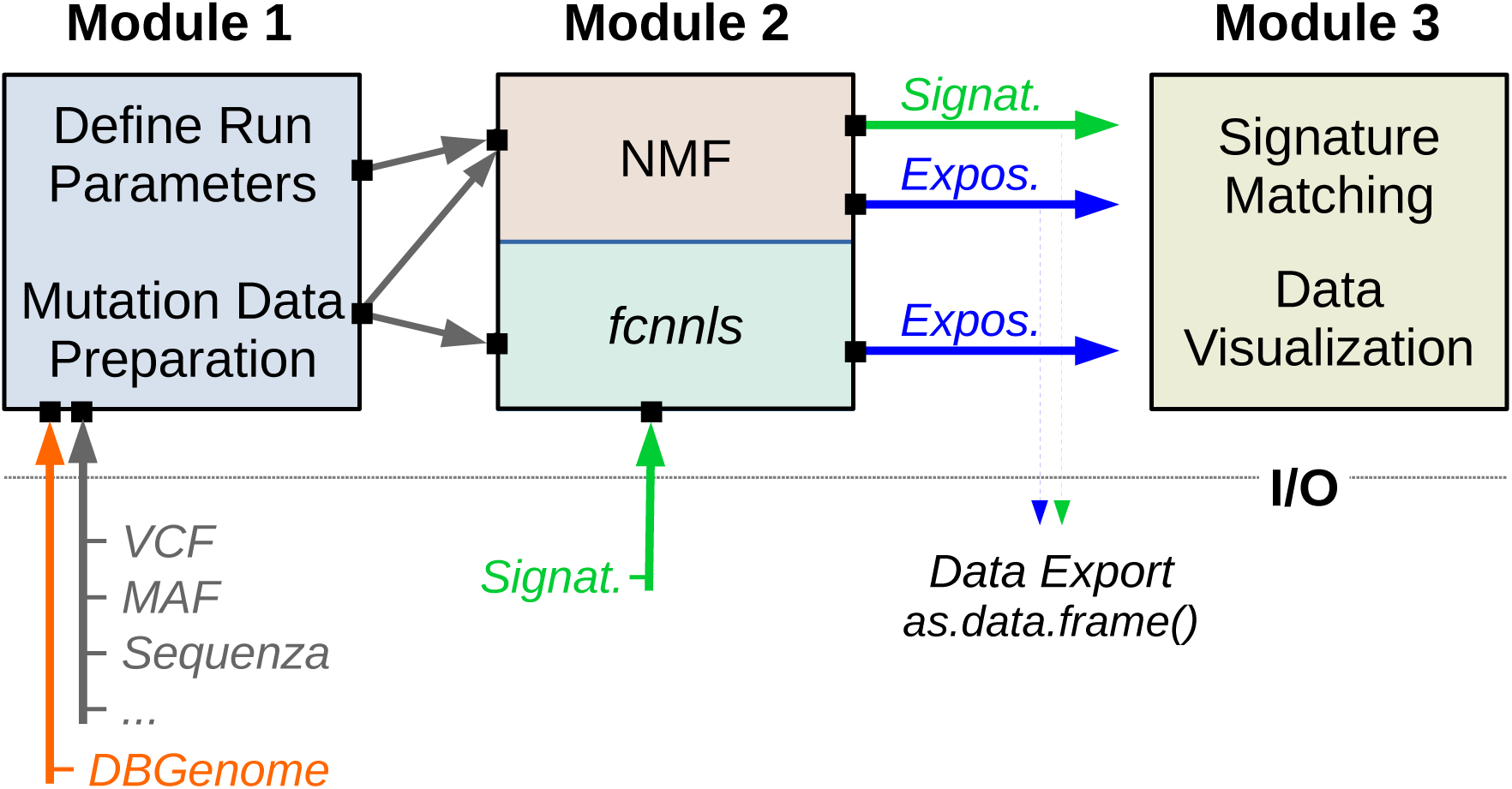
Schematic of *mutSignatures* Modules. Diagram summarizing the three modules of the *mutSignatures* framework. Module 1 is aimed at importing and preparing mutation data from VCF files or other sources. A *DBGenome* object is required for computing mutation types. Analytic parameters are set before running NMF. Module 2 is aimed at extracting mutational signatures by NMF, or computing signature exposures via the *fcnnls* function. Module 3 includes functions for comparing mutational signatures and data visualization. A summary of Input/Output (I/O) objects is shown.

### Data Import and Preparation

The *mutSignatures* framework can import DNA mutation data from multiple sources. VCF files, which are typically used to record DNA variants, can be imported individually or in batch. MAF files, used by The Cancer Genome Atlas (TCGA) to store cancer mutation data in tabular format, can be easily read in R and analyzed via *mutSignatures*. DNA variant data from *cBioPortal* (27) can be programmatically accessed using R packages such as *TCGAretriever* (https://CRAN.R-project.org/package=TCGAretriever), and then analyzed by *mutSignatures.* The *mutSignatures* framework can also import and process mutations revealed through the *Sequenza* pipeline (28). After single nucleotide variants are imported, their genomic location is used to extract the *n*-nucleotide (by default, *n*=3) context (centered on the mutated position) from a *BSgenome* reference assembly (for example, hg19 https://doi.org/doi:10.18129/B9.bioc.BSgenome.Hsapiens.UCSC.hg19). Our framework allows import and analysis of mutation data aligned to human as well as non-human genomes, including the mouse *mm10* assembly (21). By default, mutations types are formatted according to the style used by COSMIC and the Sanger Institute (for example, *A| C>T|A*). Reverse-complement transformation is automatically applied to display mutation types with a pyrimidine (C or T) as reference base at the mutated position. While the Sanger-derived format is adopted and recommended for consistency with previous analyses, users can opt for customized mutation dictionaries. Indeed, downstream analytic modules can accept either standard or non-standard mutation types as input. In the final data preparation step, mutation types are counted across all samples, returning a *mutationCounts* object that can be piped into the second module of the framework, or used for data visualization.

### *De novo* extraction of mutational signatures via NMF

Extraction of mutational signatures is conducted by NMF, as originally described for the WTSI framework (12), and according to the equation *V ≈ W* × *H*. Briefly, let *V* be an *m-by-n* non-negative mutation count matrix (including *m* mutation types and *n* biological samples). *V* is factorized into two non-negative matrices, *W* (*m*-by-*k* matrix) and *H* (*k*-by-*n* matrix). While *W* stores *k* mutational signatures, *H* includes signature exposures, which estimate the contribution of mutational signatures to the total number of mutations found in each sample (14).

Similar to the WTSI framework, in *mutSignatures* the NMF step is executed multiple times with the input count matrix bootstrapped according to the multinomial distribution of mutations by sample (12). The repeated bootstrapping followed by NMF is crucial to ensure identification of consistent and reliable mutational signatures (12). Therefore, this procedure was implemented in the *mutSignatures* framework as one of its essential components, unlike other analytic pipelines where bootstrapping is not performed. The reliability of *de novo* extracted signatures can be readily assessed by inspecting the silhouette plot that is automatically returned at the end of the signature extraction process (supplementary figure S1A).

In the WTSI framework, NMF is conducted according to the multiplicative update algorithm proposed by Brunet et al (13). Our software implements the same algorithm, as well as an alternative NMF method that was first described by Lin (22). Lin’s modified multiplicative update algorithm enforced convergence, had similar computational complexity per iteration as the original NMF algorithm, and was previously applied to the analysis of genomic and biomedical data (29,30). This feature was included in our software since the comparison of results from different NMF algorithms may facilitate the identification of consistent and reliable mutational signatures.

Our R package is already optimized for parallelization: *mutSignatures* can be easily deployed on high-performance computational clusters, and relies on the use of the *parallel, foreach* (https://CRAN.R-project.org/package=foreach), and *doParallel https://CRAN.R-project.org/package=doParallel)* R packages. The output is a list including a *mutationSignatures* object storing the newly extracted mutational signatures (*Results$signatures*), and a *mutSignExposures* object that includes signature exposures (*Results$exposures*).

### Optional Mutation Count Normalization

In the original WTSI framework, no count normalization is applied before NMF, and hence this approach is inherently biased toward extraction of signatures that are prominent in samples with high mutation burden. This strategy aligns with the hypothesis that a high total number of mutations in a sample may be due to many active mutational processes, and hence that sample gets a bigger weight in the mutational signature extraction. While this hypothesis is sound, there are evidences that selected mutational processes may contribute more than others to the accumulation of somatic mutations in tumors. An example is that of tumors with hyper-mutator phenotype (31). If signatures are extracted from raw mutation counts, the presence of high mutation burden samples in the dataset may prevent precise identification of mutational signatures that are relevant in a number of low-mutation burden tumor genomes. Additionally, the total number of mutations found in tumors also depends on sequencing depth and sample quality, which are important sources of variability in the analysis of clinical specimens (32). To circumvent this problem, it may be desirable to level the weight of all samples in the dataset. This can be achieved by sample-wise mutation count normalization. In *mutSignatures*, normalization is applied by setting the “*approach*” parameter to “*freq*”.

We examined the signatures extracted with or without counts normalization from the TCGA Bladder Cancer dataset (n=395; median SNV per genome, m=224, supplementary figure S1A), which includes a single tumor with hyper-mutator phenotype (case id: TCGA-DK-A6AW-01; total number of SNV, n=4455). Our analyses using normalized counts were insensitive to the hyper-mutator outlier, and returned 4 signatures matching those previously identified in bladder tumors, namely COSMIC signatures 1, 2, 5, and 13 (figure 2A, and (11)). Conversely, the results obtained using raw mutation counts as input showed a different signature, matching the mutation profile of the hyper-mutator sample (figures 2A, 2B, and supplementary figure S2), and this prevented the correct identification of other signatures, specifically signatures COSMIC 1 and 5 (figure 2B). Tumors with hyper-mutator phenotype were found in different TCGA datasets, showing consistent mutational profiles (COSMIC signature 10, figure 2C). Analysis of these datasets revealed similar disruptions in signature identification when raw mutation counts were used instead of normalized counts from the Breast Carcinoma (BRCA), the Cervical Squamous Cell Carcinoma and Endocervical Adenocarcinoma (CESC), and the Stomach Adenocarcinoma (STAD) datasets (supplementary figure S3). Nevertheless, mutation count normalization successfully identified COSMIC 10-like signatures in a number of TCGA cohorts where the hyper-mutator phenotype occurred more frequently (Rectum, READ; Colon, COAD; and Endometrial, UCEC cancer datasets, supplementary figure S3).

**Figure 2.**
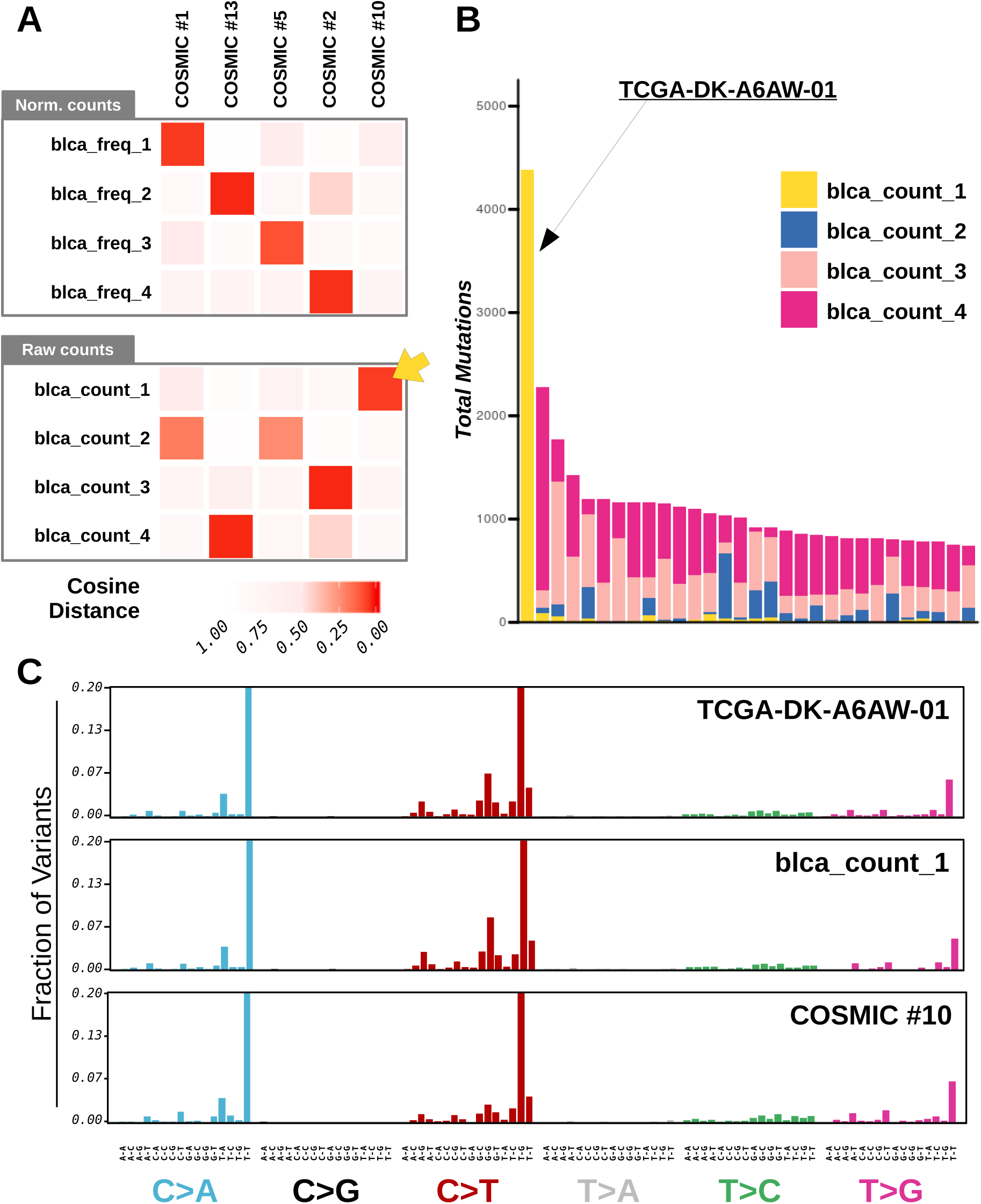
Mutational signatures identified in Bladder Cancer Genomes. (A) Heatmaps showing similarity between COSMIC signatures and mutational signatures that were *de novo* extracted from the TCGA bladder cancer dataset. Mutational signatures were identified using normalized (top heatmap) or raw (bottom heatmap) mutation counts. Cosine distances across signatures were computed, and displayed by color intensity. The yellow arrow indicates a signature that was specifically extracted when raw mutation counts were used as input. (B) Exposures to mutational signatures extracted from raw mutation counts. A limited number (n=30) of TCGA bladder cancer samples with the highest mutation burden is displayed. Each bar represents a tumor and the vertical axis denotes the number of mutations imputed to each signature (highlighted by colors). The leftmost bar of the plot (yellow bar) corresponds to the hyper-mutator sample (TCGA-DK-A6AW-01). (C) Barplots summarizing the mutational profiles of the sample (TCGA-DK-A6AW-01) and mutational signatures (blca_count_1, and COSMIC #10) corresponding to the hyper-mutator phenotype in cancer. Mutation types were grouped by SNV.

### Deconvolution of mutation counts against known mutational signatures

Computing exposures when mutational signatures are known means solving the *V ≈ W* × *H* equation when both *V* and *W* are known and *H* is unknown. Our framework solves this nonnegative least square linear problem via a custom implementation of the fast combinatorial strategy proposed by Van Benthem (33). Imputed signature exposures are returned as a *mutSignExposures* object. Removal of under-represented signatures is not automatically applied. The *deconstructSigs* R package (26) is dedicated to this kind of analysis, and returned overlapping results when compared to our method (supplementary figure S4), with our approach being about 50 times faster than *deconstructSigs*. Recently, the strategy of using our *mutSignatures* package for *de novo* signature extraction alongside with *deconstructSigs* for mutation counts deconvolution has been successfully implemented (34).

## RESULTS

### Extraction of mutational signatures from smoking-related cancers

A link between DNA mutational signatures and tobacco consumption was reported before (16,17,35), showing that tumors from smokers had higher mutation burden compared to non-smokers, and that prevalent mutational signatures in smoking-related cancers were COSMIC signatures 4, 5 (16,35), as well as the APOBEC-associated signatures (COSMIC signatures 2 and 13) (35-37). Here we used the *mutSignatures* framework to extract tri- and tetra-nucleotide mutational signatures from the lung adenocarcinoma (*LUAD*) TCGA dataset, and analyzed correlations with other molecular or clinical parameters. Samples with at least 50 total SNV (supplementary figure S5A) per genome and including information about survival and tobacco smoking history were analyzed (figure 3A). We found that genomes of current or reformed smokers had significant (t-test p-val ≤ 2.0e-13) accumulation of mutations compared to life-long non-smokers (figure 3B). Stage I tumors showed statistically (log-rank p-val ≤ 6.5e-05) better survival compared to higher tumor stages (figure 3C). On the contrary, smoking status was not indicative of clinical outcomes (supplementary figure S5B). Tri- and tetra-nucleotide signatures were extracted from the 418 genomes meeting the inclusion criteria.

**Figure 3.**
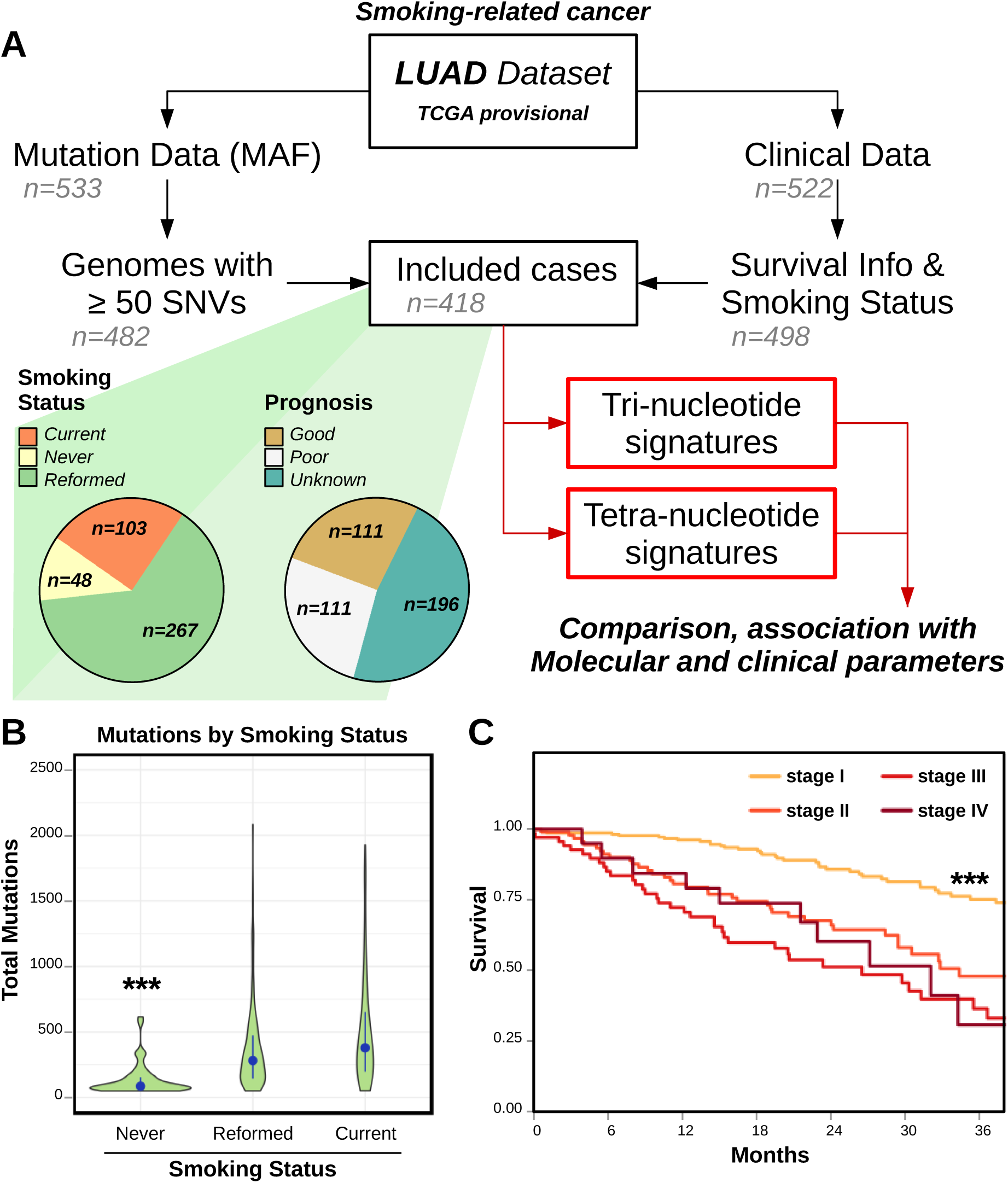
Mutational landscape of Lung Adenocarcinoma. (A) Diagram summarizing the sample inclusion criteria applied for the analysis of the LUAD TCGA dataset. Patients with both survival and tobacco consumption information, and including at least 50 SNV in their genome were analyzed (n=418). Samples were used for tri- and tetra-nucleotide signature extraction. Pie charts summarize the distribution of smoking status and prognosis in the included patients. (B) Violin plot showing the distribution of total number of mutations detected in LUAD cancer genomes according to the patients’ smoking status. Blue dots indicate the median values; blue segments indicate the range spanning from the first to the third quartile. Groups were compared by t-test. (C) Plot comparing survival of LUAD cancer patients according to tumor stage (I to IV). Groups were compared by log-rank test. Three asterisks (***) indicate p-value less than 1e-4 for the labelled group compared to all others.

### Comparison between tri- and tetra-nucleotide mutational signatures

Tri-nucleotide mutational signatures extracted from the *LUAD* TCGA dataset matched COSMIC signatures 1, 2, 4, and 5 (figure 4A, and supplementary figure S6, previously identified in lung cancer genomes (11). Next, we examined tetra-nucleotide signatures, which were obtained from DNA mutation types including information about the nucleotide at the 5’-end of the standard tri-nucleotide mutations. To allow comparison with standard mutational signatures, we aggregated frequencies of tetra-nucleotide DNA variants corresponding the same tri-nucleotide mutation type. This operation returned a list of simplified tetra-nucleotide signatures that overlapped with the tri-nucleotide mutational signatures derived before (figures 4B, 4C, and supplementary figure S6). The close similarity between signatures extracted via either method demonstrated the reliability of results obtained using our analytic framework and the context-specificity of mutational signatures. A closer inspection of tetra-nucleotide signatures confirmed the sensitivity of mutations to their flanking DNA sequences, including not only the immediate neighboring bases, but also the second base at the 5’-end of selected SNV. For example, signature *luad_tetra_B* featured a striking preference for cytosine upstream of C|C>A|N, as well as of T|C>A|G mutations (figure 4C, supplementary figure S6), similar to previous reports (17). Therefore, our observations supported that the study of extended mutation types (such as tetra-nucleotide mutations) could carry more complete information and provide insights in the biology underlying DNA mutagenesis in cancer.

**Figure 4.**
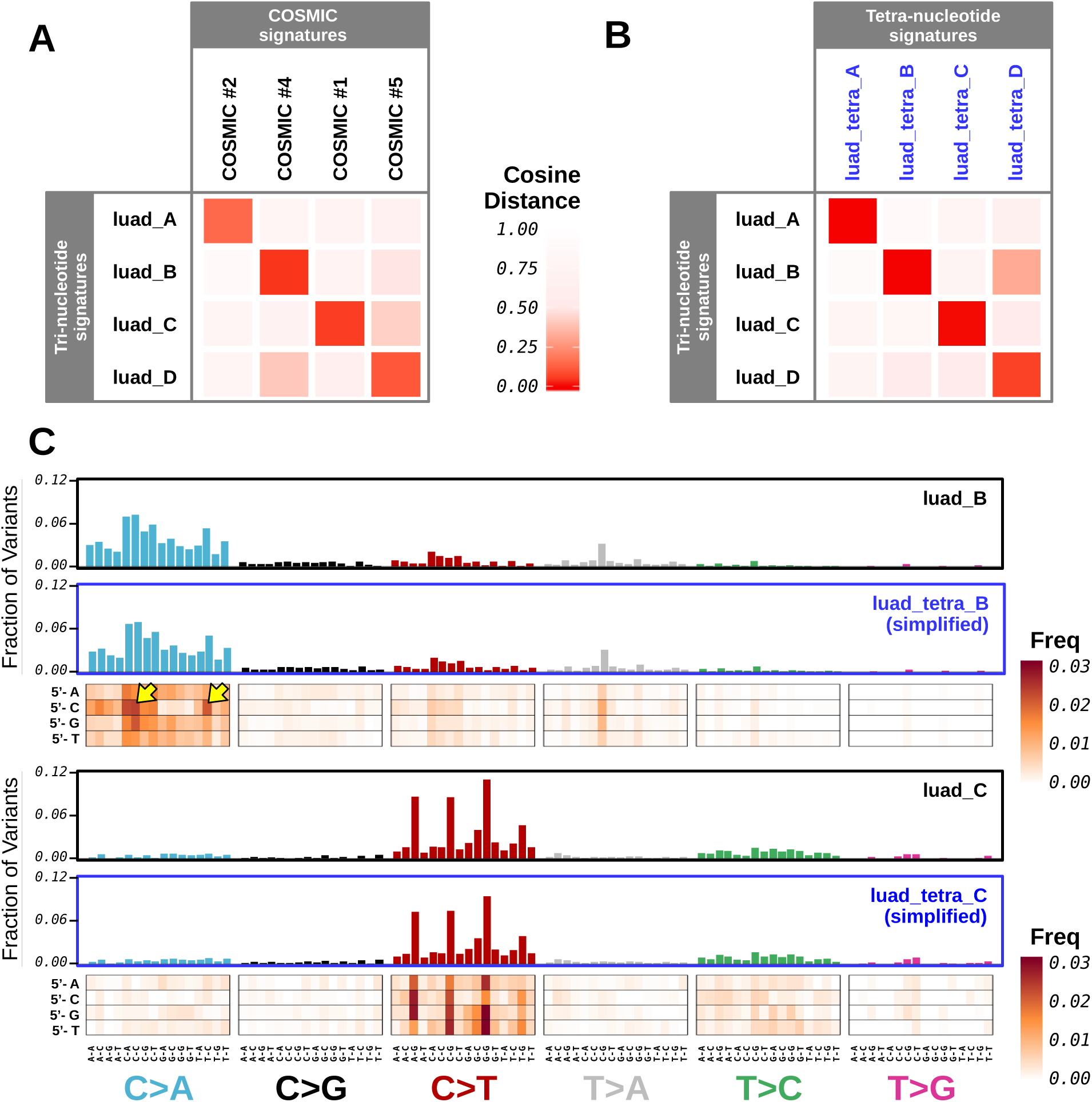
Analysis of tri- and tetra-nucleotide mutational signatures extracted from the LUAD TCGA dataset. (A) Heatmap examining similarity between COSMIC signatures, and tri-nucleotide mutational signatures that were *de novo* extracted from the LUAD TCGA dataset. (B) Heatmap comparing tri- and tetra-nucleotide mutational signatures that were *de novo* extracted from the LUAD TCGA dataset. Tetra-nucleotide signatures were simplified to the corresponding tri-nucleotide signatures by mutation type binning. Color intensity tracks with the value of cosine distance. (C) Barplots and heatmaps summarizing the mutational profiles of tri- and tetra-nucleotide mutational signatures (top: *luad_B* and *luad_tetra_B*; bottom: *luad_C*, and *luad_tetra_C*). Heatmaps are visual representations of the tetra-nucleotide mutational signatures, where tri-nucleotide mutation types are shown on the x-axis, and the extra 5’-end nucleotides are shown on the y-axis. Box color intensity tracks with mutation type frequency. Tetra-signature simplification can be summarized as the result of column-wise aggregation of tetra-nucleotide mutation frequencies as shown in the heatmaps. Simplification returned vectors of tri-nucleotide mutation type frequency that are displayed as barplots.

### Mutational signature Exposures in LUAD TCGA genomes

We analyzed the tri-nucleotide mutational signature exposures across lung cancer samples. Signature exposures indicate how many mutations are the consequence of each mutational signature in each sample (figure 5A). Analysis of signature exposures revealed two groups in the data: i) tumors enriched in *luad_B* signature, usually having high mutation burden (group 1); and ii) tumors depleted in *luad_B* signature, usually featuring low total number of DNA mutations (group 2). Analysis of relative exposures (exposures normalized by total number of mutations in the genome) showed that the *luad_C* signature was enriched in group 2 samples (figure 5A).

**Figure 5.**
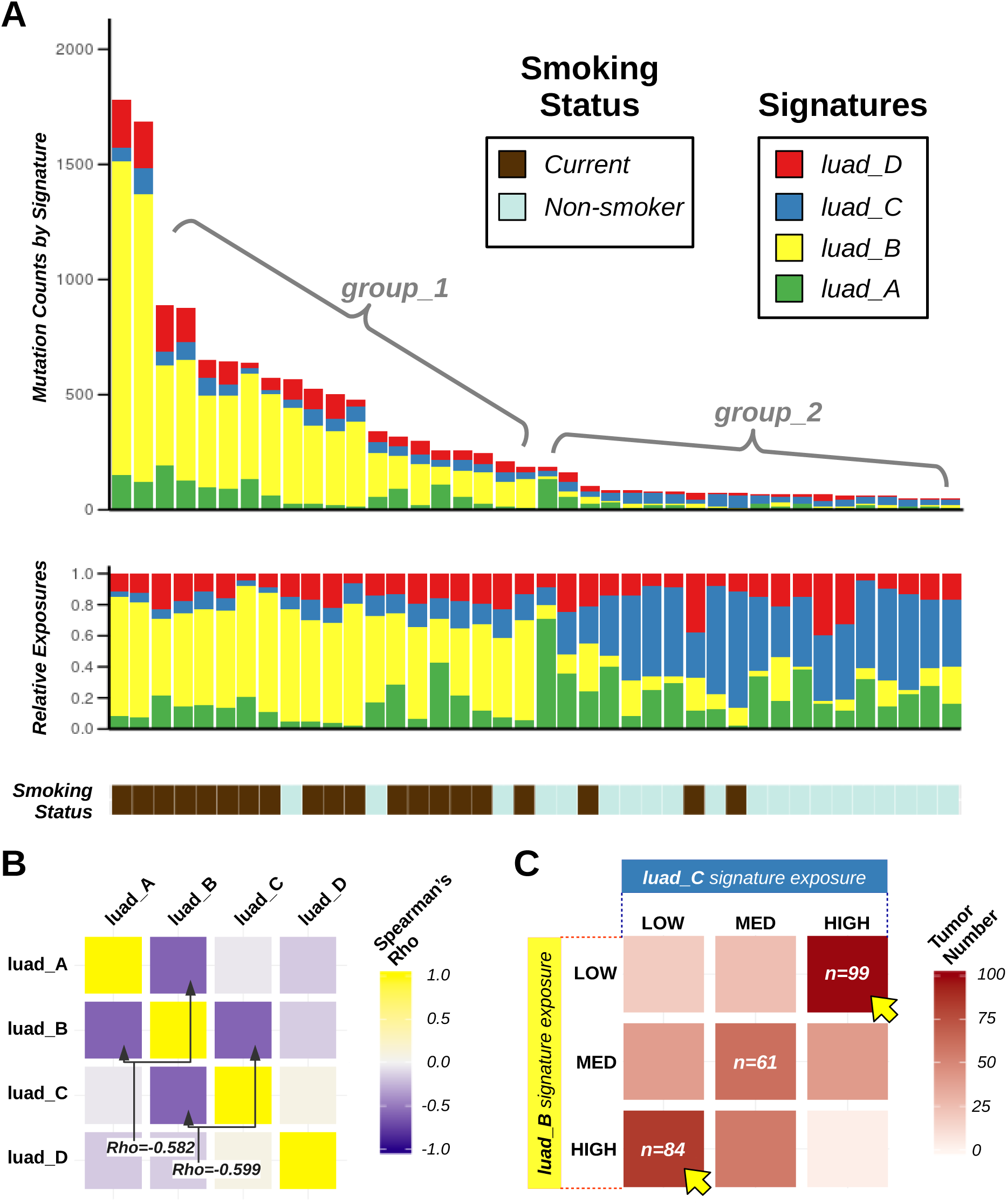
Signature Exposures in smokers and non-smokers affected by lung adenocarcinoma. A) Exposures to mutational signatures that were *de novo* extracted from LUAD TCGA. A limited number (n=40, including 20 random genomes from smokers and 20 random genomes from life-long non-smokers) of lung cancer samples are displayed. Each bar represents a tumor and the vertical axis denotes the total (top barplot) or the relative (central barplot) number of mutations imputed to each signature (highlighted by colors). The patient smoking status key is shown below the barplots. B) Heatmap showing *Spearman* correlation coefficients (Rho) across signature exposures in the Lung Adenocarcinoma dataset. Exposures to standard tri-nucleotide mutational signatures were analyzed. Yellow boxes correspond to positive correlations; blue boxes indicate pairs of signatures that are inversely correlated. C) Heatmap highlighting the distribution of exposures to *luad_B* (y-axis) and *luad_C* (x-axis) signatures in LUAD TCGA genomes. Exposures to both signatures were tertile-discretized (low, medium, and high), and then orthogonally analyzed. Tumors belonging to each of the 9 possible groups were counted. Color intensity tracks with number of patients. Yellow arrows indicate the two groups with the highest patient count, which corresponded to tumors with high exposure to one signature and low exposure to the other.

We computed *Spearman* correlation between relative signature exposures (figure 5B), and confirmed our previous observations. The pairs of signatures with the lowest Spearman’s coefficient were signatures A and B (Rho=-0.582), and signatures B and C (Rho=-0.599), while signatures A and C were uncorrelated (Rho=-0.047). Negative correlations among mutational signatures were anticipated because of the constraint that relative exposures had to sum up to unity, but the observed Rho values were significantly lower compared to those expected according to Monte Carlo simulations (p<0.005, supplementary figure S7A). In addition, we quantile-discretized and examined relative exposures to signatures B and C, and found that tumors were more likely to have high contribution of one or the other signature rather than intermediate exposure of both of them (figure 5B).

Notably, these two signatures matched signatures COSMIC 4 and 1, respectively (figure 4A). COSMIC 4 was proposed to originate after the activity of cigarette smoke carcinogens, while COSMIC 1 was associated to spontaneous deamination of 5-methylcytosine. Our observations suggested that these two signatures and the corresponding mutational processes had a tendency to occur in mutual exclusive fashion in lung adenocarcinoma.

### Mutational signatures and clinical parameters

We further analyzed mutational signatures and their associations with molecular and clinical parameters. First, we compared mutational signatures and mutation burden. In agreement with what observed before, we found that signature *luad_B* was significantly enriched in high mutation burden genomes (Kendall’s rank correlation test, tau = 0.4563, p-val < 2.2e-16, figure 6A), and that the relative contribution of signature *luad_C* was higher in low mutation burden samples (Kendall’s rank correlation test, tau = -0.6240, p-val < 2.2e-16, figure 6B).

Next, we tested whether mutational signatures were prognostic of patient clinical parameters. We could not find any correlation between mutational signatures and overall patient survival (supplementary figure S7B). However, we tested whether signatures *luad_B* and *luad_C* were significantly correlated with other clinical features, especially patient smoking status. Our analyses revealed that exposures to signature *luad_B* were increased (t-test, p-val < 3.4e-10) in tumors from smokers (both current and reformed, figure 5C). Conversely, relative exposures to signature *luad_C* was increased in life-long non-smokers (t-test, p-val < 6.7e-6, figure 5D). To validate our conclusions, we examined the association between *luad_B* and *luad_C* mutational signatures and clinical features in a different smoking-related cancer dataset. We analyzed the Head and Neck Squamous Cell Carcinoma (HNSC) because the mutational signatures identified in this dataset using the *WTSI MATLAB* framework were similar to those detected by COSMIC in lung adenocarcinoma. We deconvoluted mutation catalogs from the HNSC TCGA dataset (n=511) against the four signatures extracted from LUAD TCGA (*luad_A, luad_B, luad_C*, and *luad_D*). Next, we assessed the association between smoking status and relative exposures. In agreement with our observations, we found that signature *luad_B* was significantly higher in genomes of smoking HNSC patients (figure 6E; t-test, non-smokers vs. smokers, p-val<3.4e-06), while relative exposures to signature *luad_C* were higher in head and neck tumors from non-smoking patients (figure 6F; t-test, non-smokers vs. smokers, p-val<2.6e-05).

**Figure 6.**
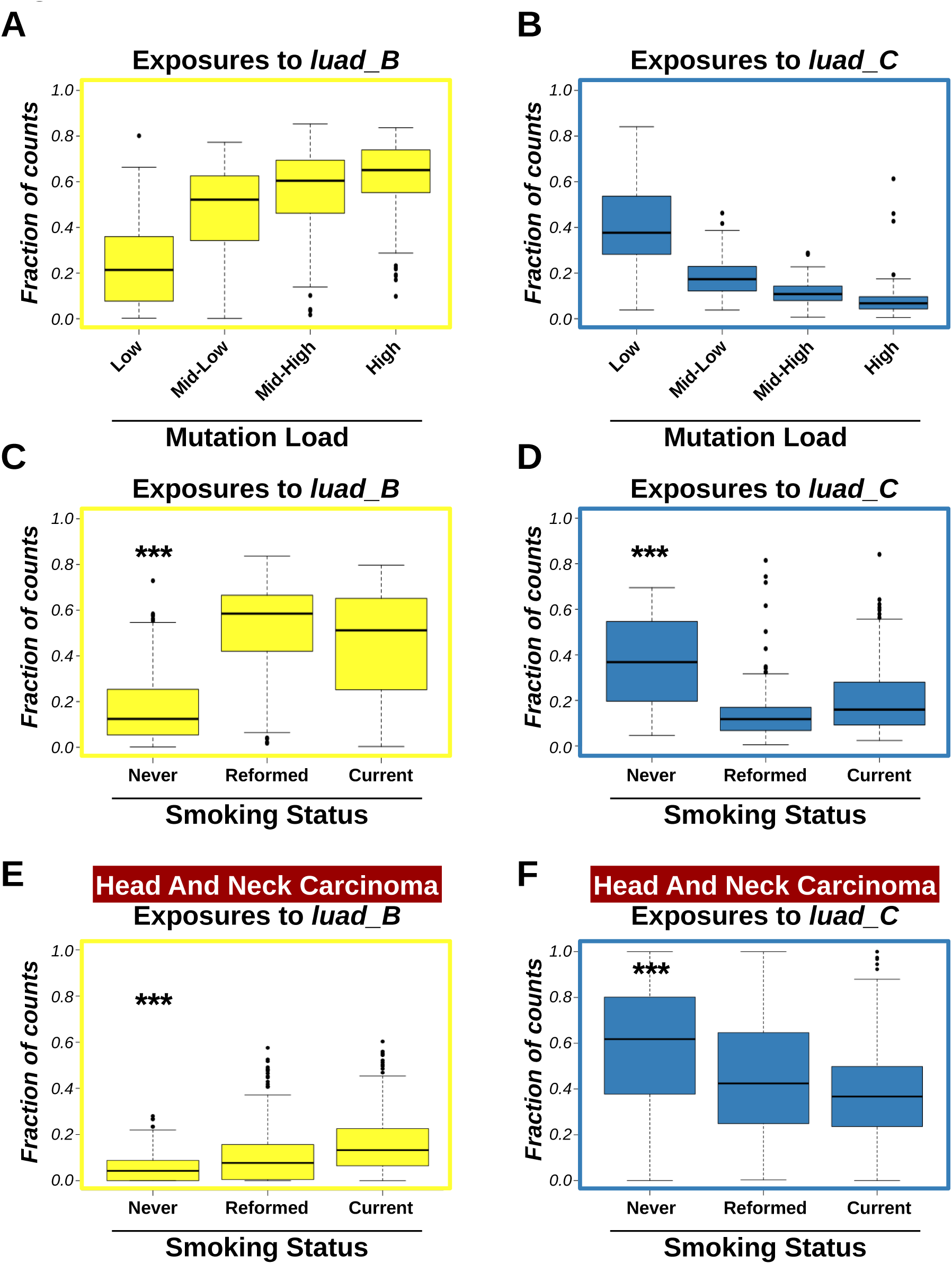
Correlation among mutational signatures, mutation burden, and smoking status in lung adenocarcinomas. A, B) Boxplots showing relative exposure to signatures *luad_B* (A) and *luad_C* (B) according to discretized mutation burden. Mutation burden was quartile-discretized. Correlation between relative exposures and binned mutation burden was computed by Kendall’s rank correlation test. Kendall’s coefficients (tau) were tau=0.4563 (p-val < 2.2e-16) for *luad_B* signature (A), and tau=-0.6240 (p-val < 2.2e-16) for *luad_C* signature (B). C, D) Boxplots showing relative exposure to signatures *luad_B* (C) and *luad_C* (D) in LUAD genomes according to patient smoking status. Groups were compared by *t-test*. E, F) Boxplots showing relative exposure to signatures *luad_B* (E) and *luad_C* (F) in HNSC genomes according to patient smoking status. Groups were compared by *t-test*. Three asterisks (***) indicate p-val less than 1e-4.

Our results showed that *mutSignatures* supported the characterization of genetic instability mechanisms active in lung adenocarcinoma, and revealed mutational signatures that were strongly associated with specific molecular and clinical parameters, such as mutation burden, and patient smoking history. Likewise, similar analyses may enable prediction of other signature-associated clinical parameters, for example response to selected anticancer therapies, and ultimately support gathering insights into tumor biology and treatment.

## DISCUSSION

Identifying the molecular mechanisms driving tumor initiation and progression is crucial in cancer research and therapeutics. The study of DNA mutational signatures is an emerging area of cancer genomics that can help understanding what mechanisms are responsible for the accumulation of somatic mutations found in tumors. Here, we introduced *mutSignatures*, a software supporting extraction and analysis of DNA mutational signatures. Our framework is written in R (https://www.r-project.org/), a free statistical programming environment, and aligns to the standards set by the WTSI MATLAB framework by Alexandrov et al (12). Moreover, our software includes tools for mutation data import and preparation, mutational signature extraction and analysis via non-negative matrix factorization, and data visualization. Compared to the original WTSI framework and other available software, *mutSignatures* includes new functionalities that address some of the current methodological limitations. For example, our framework is compatible with non-human genomes, can extract and analyze non-standard mutation types, and enables built-in sample-wise mutation count normalization. Moreover, *mutSignatures* can be easily streamlined with existing R libraries and R-based genomic analytic pipelines.

Here, we used *mutSignatures* to extract and analyze mutational signatures from TCGA lung adenocarcinoma genomes and other datasets. We successfully identified mutational signatures matching those previously reported by COSMIC in the same types of cancer. For the first time, we extracted tri- and tetra-nucleotide mutational signatures using the same algorithm. Our characterization revealed a great similarity between signatures obtained using standard or non-standard mutation types, confirming the reliability of the analytical approach implemented in our R framework, as well as the nucleotide-context specificity of mutational signatures. Our results showed that DNA mutations are highly sensitive to their nucleotide context, which is not solely limited to the immediate flanking bases but extends further. This provides rationale for the study of non-standard extended (more than 3 nucleotides) mutation types, a kind of analysis that is supported by *mutSignatures*.

Finally, we analyzed correlations between mutational signatures found in lung adenocarcinoma samples, and other clinical and molecular features. We identified two signatures, namely *luad_B* and *luad_C*, which were inversely correlated. Signature *luad_B* was increased in tumors from smokers and correlated with high mutation burden. Conversely, signature *luad_C* was enriched in tumors from life-long non-smokers, and correlated with low mutation burden. These two signatures may be the consequence of mutually-exclusive mutational processes resulting in the incorporation of DNA mutations in lung cancer cells from smoking and non-smoking patients, respectively. Similarly, mutational signature analyses could reveal correlations with other molecular or clinical parameters, such as expected clinical course, or patient response to specific anti-cancer drugs.

In conclusion, we presented *mutSignatures*, an R package for analysis of mutational signatures. Our software can be used for the identification of mutational determinants of cancer, supports the analysis of signature-associated molecular and clinical features, and has the potential of revealing insights into tumor biology and treatment.

## AVAILABILITY

The latest version of *mutSignatures* (version 2.0.1) is available on CRAN or at the following URL: https://github.com/dami82/mutSignatures.

## ACKNOWLEDGEMENT

DF, JJM designed the research project. DF, YY acquired and prepared the data. DF developed software, wrote R extension, performed data analyses. DF and VV contributed to data visualization. DF, JJM, VV, SC wrote and reviewed the manuscript.

