## Supplementary Material for "MutSignatures: An R Package for Extraction and Analysis of Cancer Mutational Signatures"

### Supplementary Figure S1

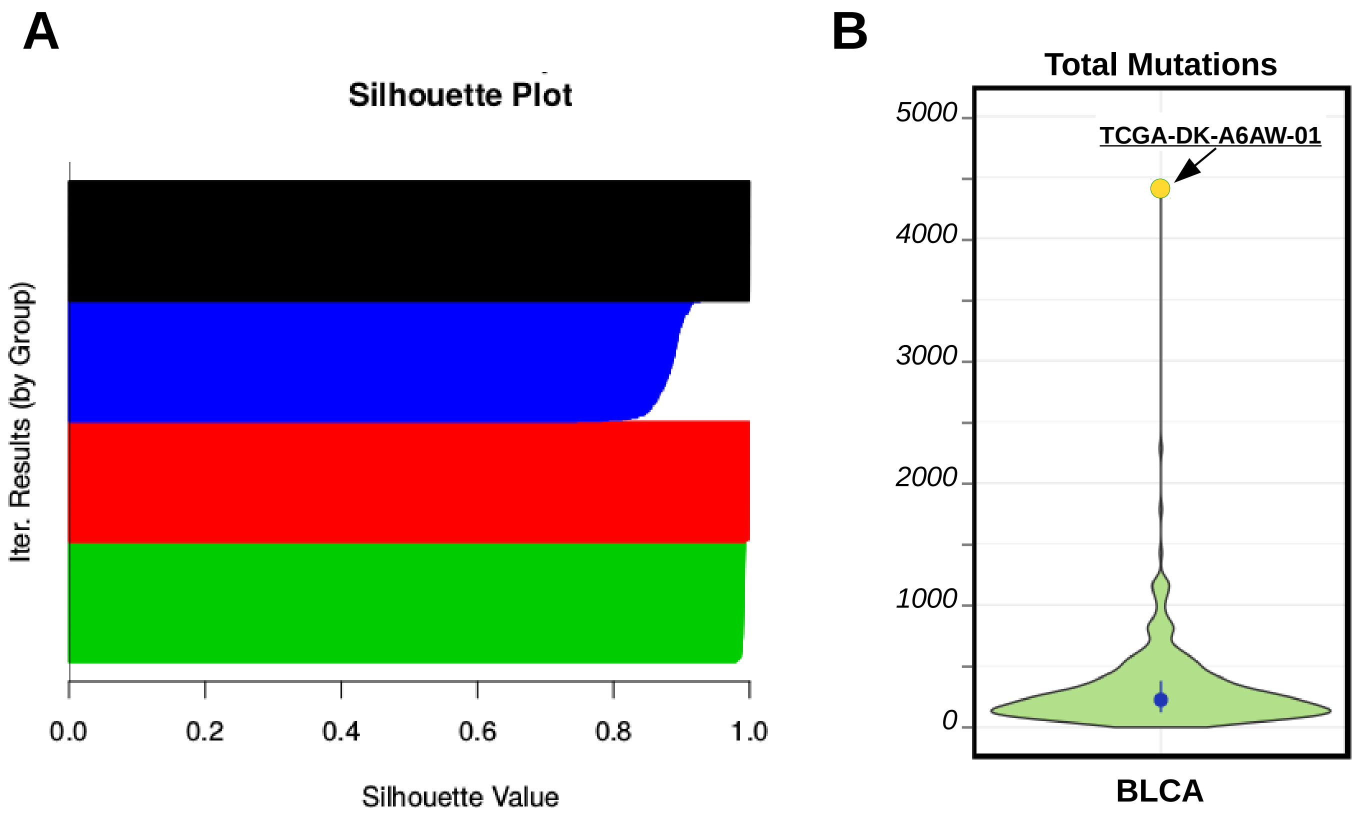


**Supplementary Figure S1.** Mutational landscape of TCGA Bladder Cancer. A) Silhouette plot (horizontal barplot) returned at the end of a *de novo* mutational signature extraction. Each of the *k* (here, *k*=4) signatures extracted during each iteration (here, n=500) is assigned to one of the *k* possible groups (identified by colors) based on similarity to group centroids. Distances to centroids are then plotted (horizontal bars). If all the signatures that are assigned to the same group are very similar, signature identification is solid and reliable. In the silhouette plot, this results in bars close to the value 1 (as shown in the plot). Barplot groups showing poor silhouette values (values close to 0 or negative values) are suggestive of unreliable signature extraction (usually, this means that too many signatures were extracted). B) Violin plot showing the distribution of mutation burden in the BLCA TCGA dataset. The yellow point corresponds to the hyper-mutator sample (id: TCGA-DK-A6AW- 01). The y-axis indicates number of SNV detected per genome.

### Supplementary Figure S2

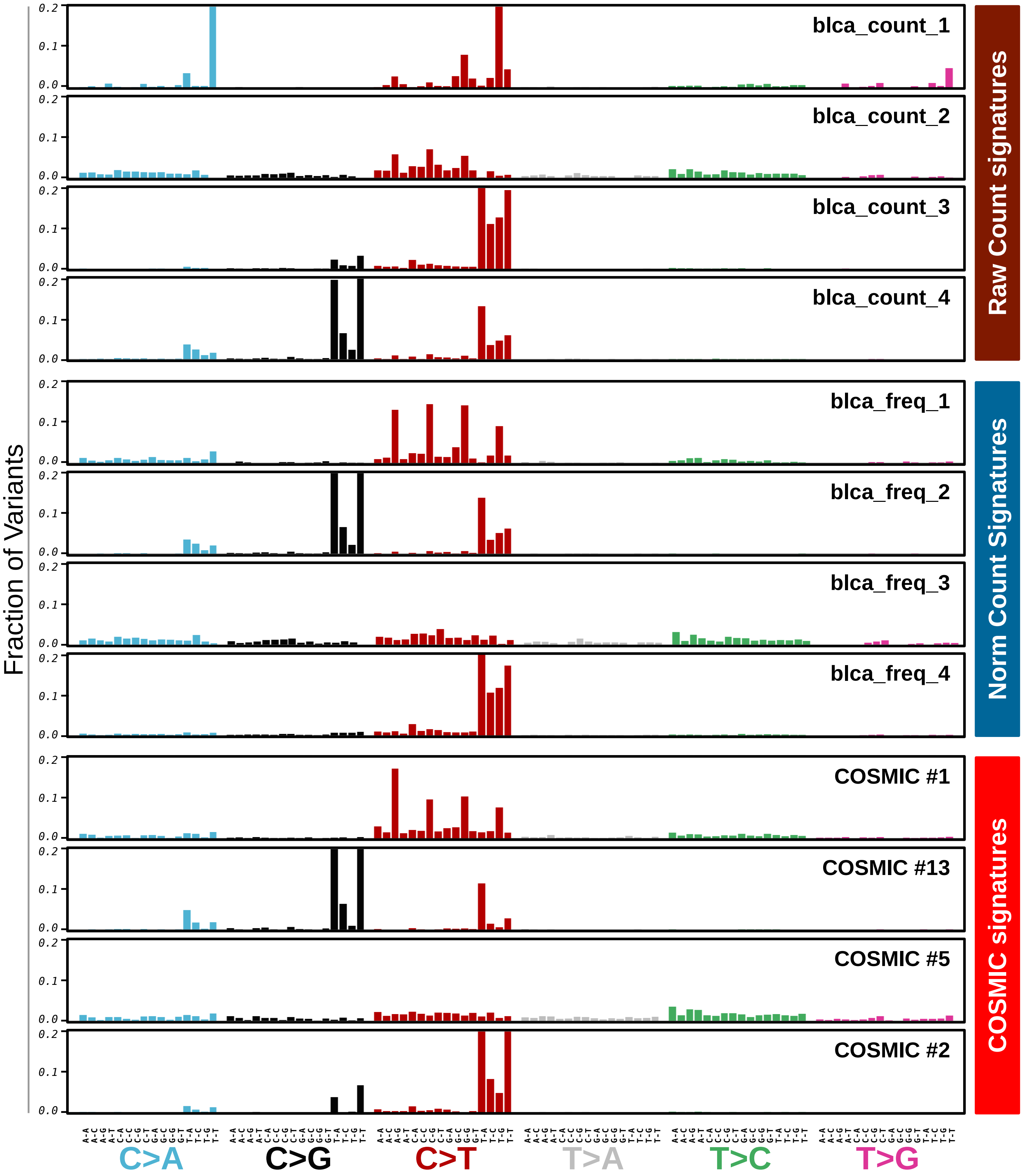


**Supplementary Figure S2.** Mutational signatures extracted from BLCA TCGA. Barplots summarizing the mutational profiles of mutational signatures extracted from TCGA bladder cancer genomes, and the corresponding COSMIC signatures are displayed. Single nucleotide variants were grouped by the tri-nucleotide mutation type.

### Supplementary Figure S3

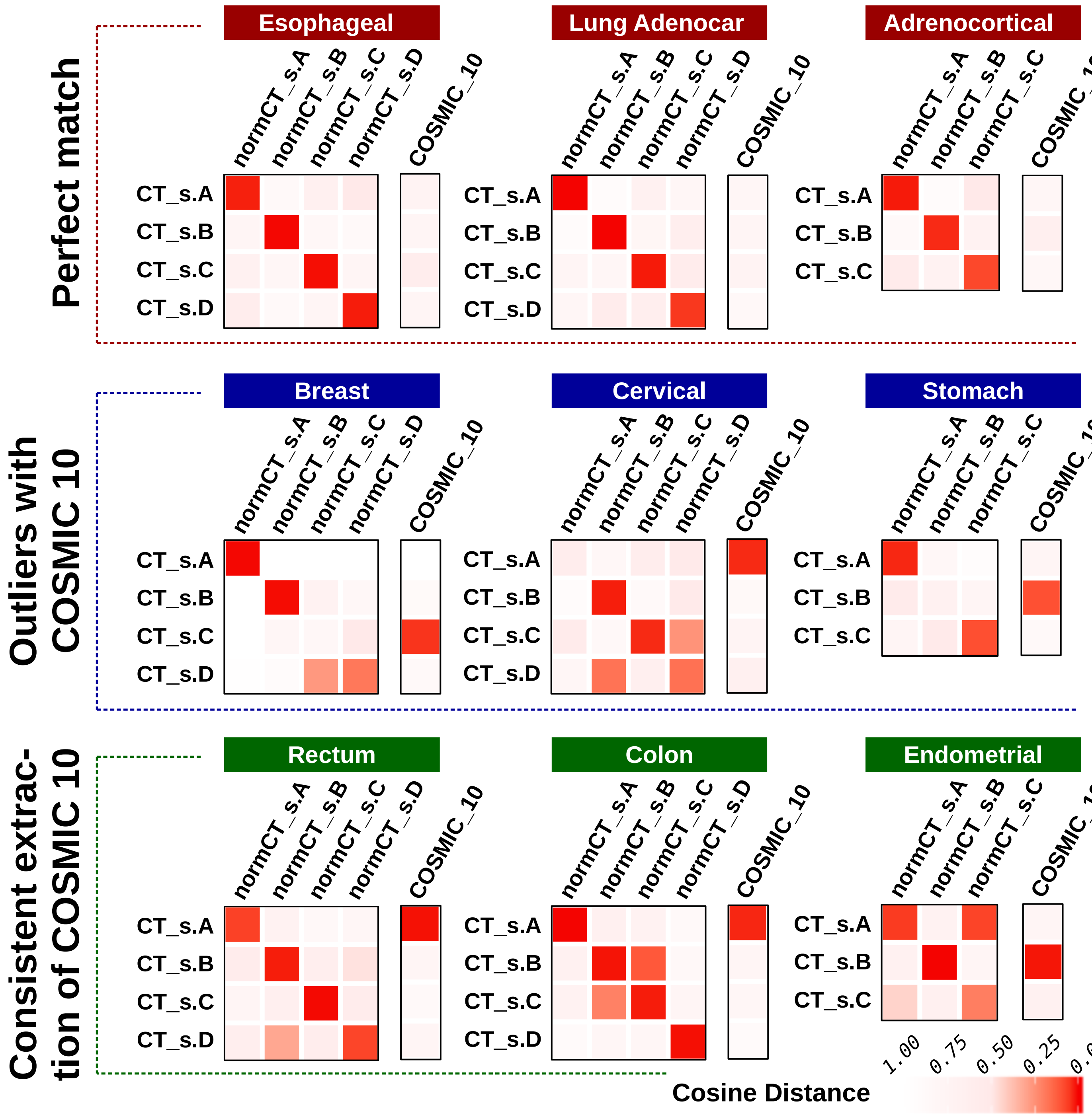


**Supplementary Figure S3.** Comparison between mutational signatures extracted using raw or normalized mutation counts. DNA Variants from 9 TCGA datasets (ESCA, LUAD, ACC, BRCA, CESC, STAD, READ, COAD, UCEC) were imported and analyzed via *mutSignatures*. Mutational signatures were extracted from genomes with at least 50 SNV. Signatures from raw mutation counts (CT_s.A-to-D, y-axis) were compared to COSMIC 10 signature, as well as signatures extracted after application of count normalization (normCT_s.A-to-D, x-axis). Cosine distance between signatures was computed and displayed by heatmaps (red boxes, high similarity; white boxes, high dissimilarity). Three distinct scenarios were observed.

1) Perfect Match (top): consistent signatures were extracted by either method, and COSMIC 10-like signatures were not identified; 2) Outliers with COSMIC 10 (middle): one of the signatures extracted using raw counts matched COSMIC 10 signature, but this signature was not detected upon count normalization; 3) Consistent Extraction of COSMIC 10 (bottom): either approach returned a COSMIC 10-like signature, suggesting that samples with hyper-mutator phenotype were common in the dataset.

### Supplementary Figure S4

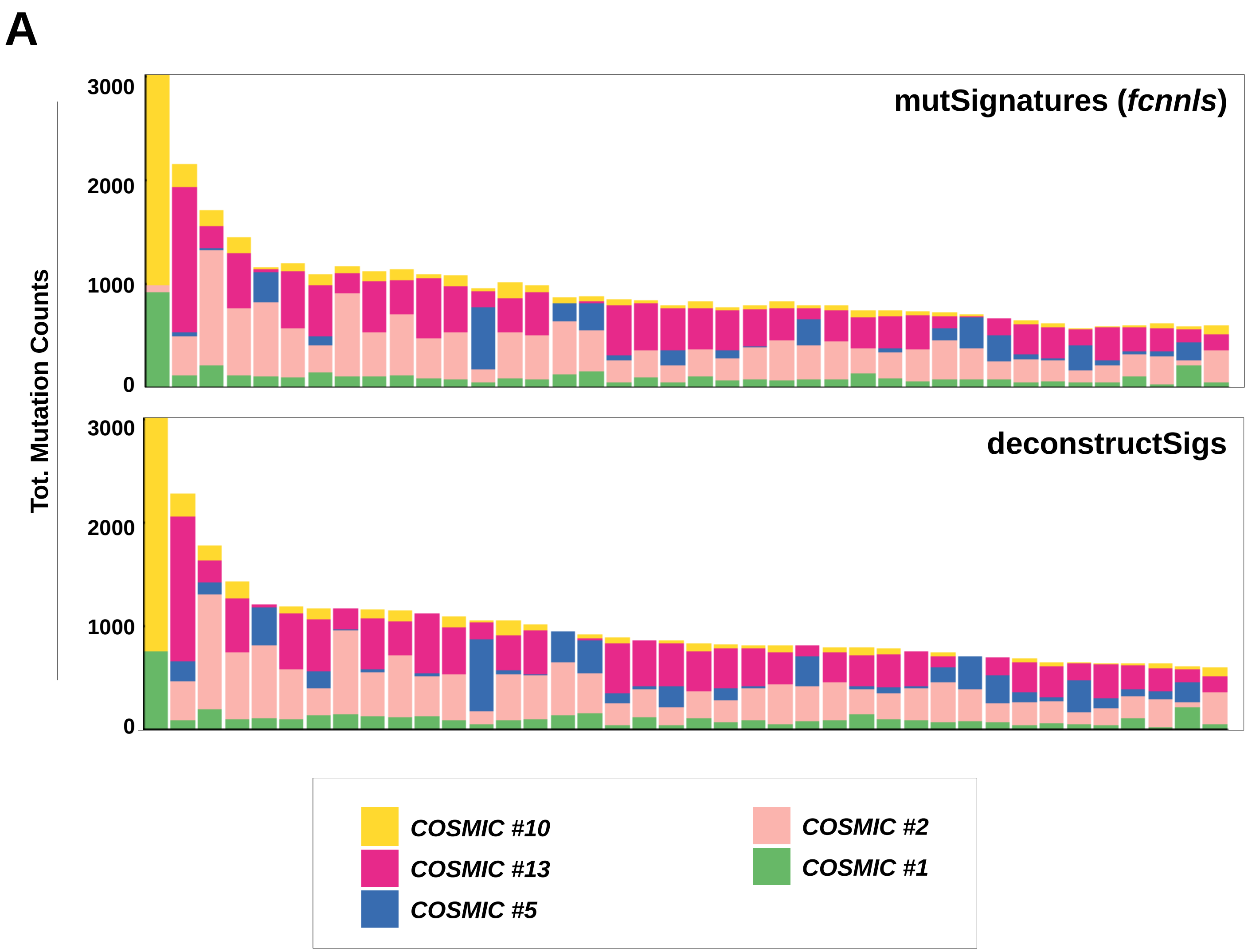


**Supplementary Figure S4.** Deconvolution of mutation counts against COSMIC mutational signatures. BLCA TCGA mutation counts were deconvoluted against COSMIC signatures 1, 2, 5, 10, and 13 using *mutSignatures* or *deconstructSigs*. The top 40 samples according to mutation load were included in the barplot. Each bar represents a tumor and the vertical axis denotes the number of mutations imputed to each signature.

### Supplementary Figure S5

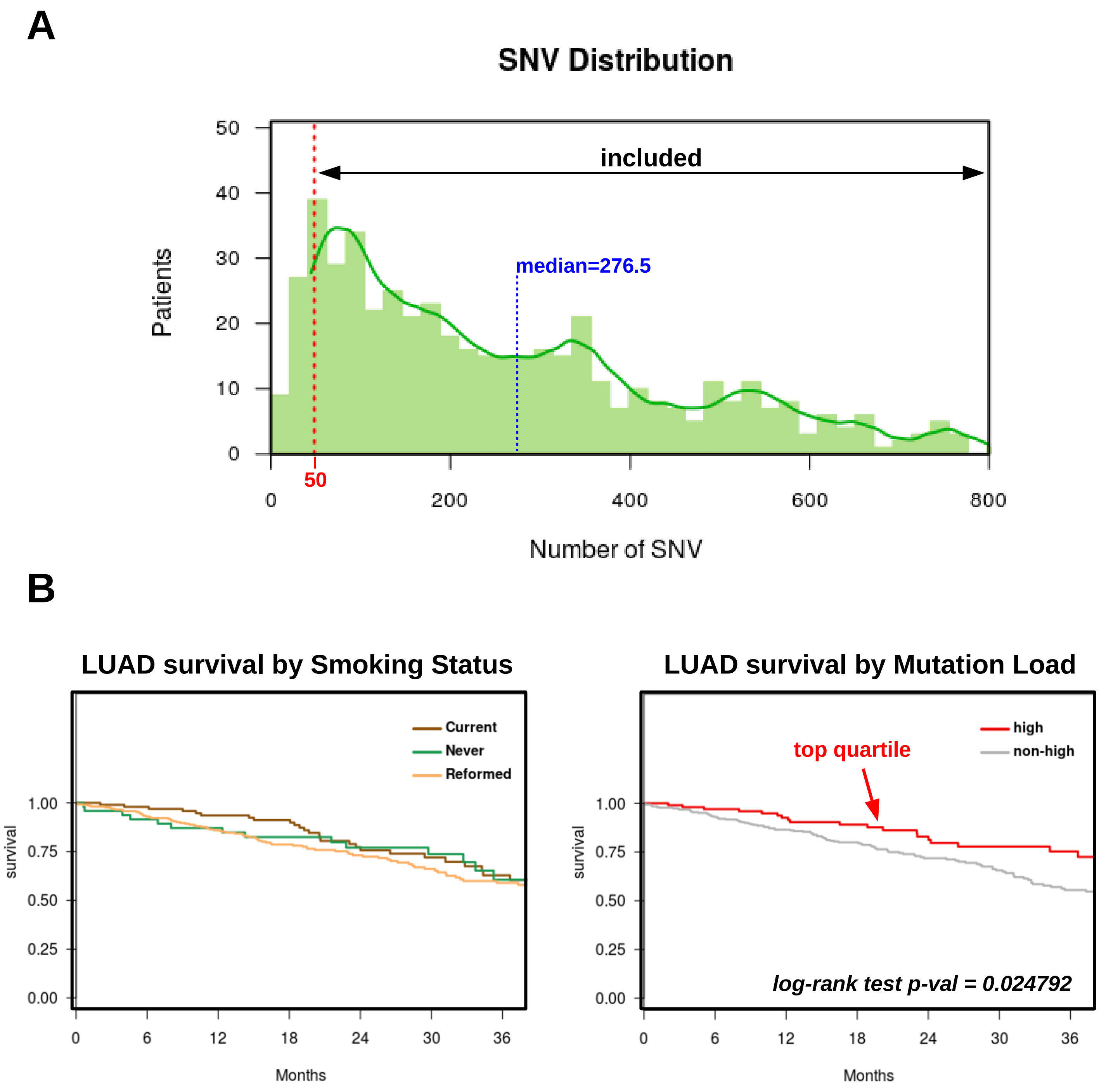


**Supplementary Figure S5.** Mutational landscape of the LUAD TCGA dataset. A) Histogram showing the distribution of total SNV mutation in the samples of LUAD TCGA. The y-axis indicates the number of patients, the x-axis shows the number of SNV per genome. The median SNV per genome was 276.5 (blue segment). The red dotted line show the inclusion threshold (samples with less than 50 SNV per genomes were excluded from the analysis). B) Patient Survival was analyzed according to smoking status (left plot) or mutation load (right). Patients with high mutation burden (top quartile) had slightly better survival than other patients (right). Groups were analyzed by log-rank test (p-val=0.025).

### Supplementary Figure S6

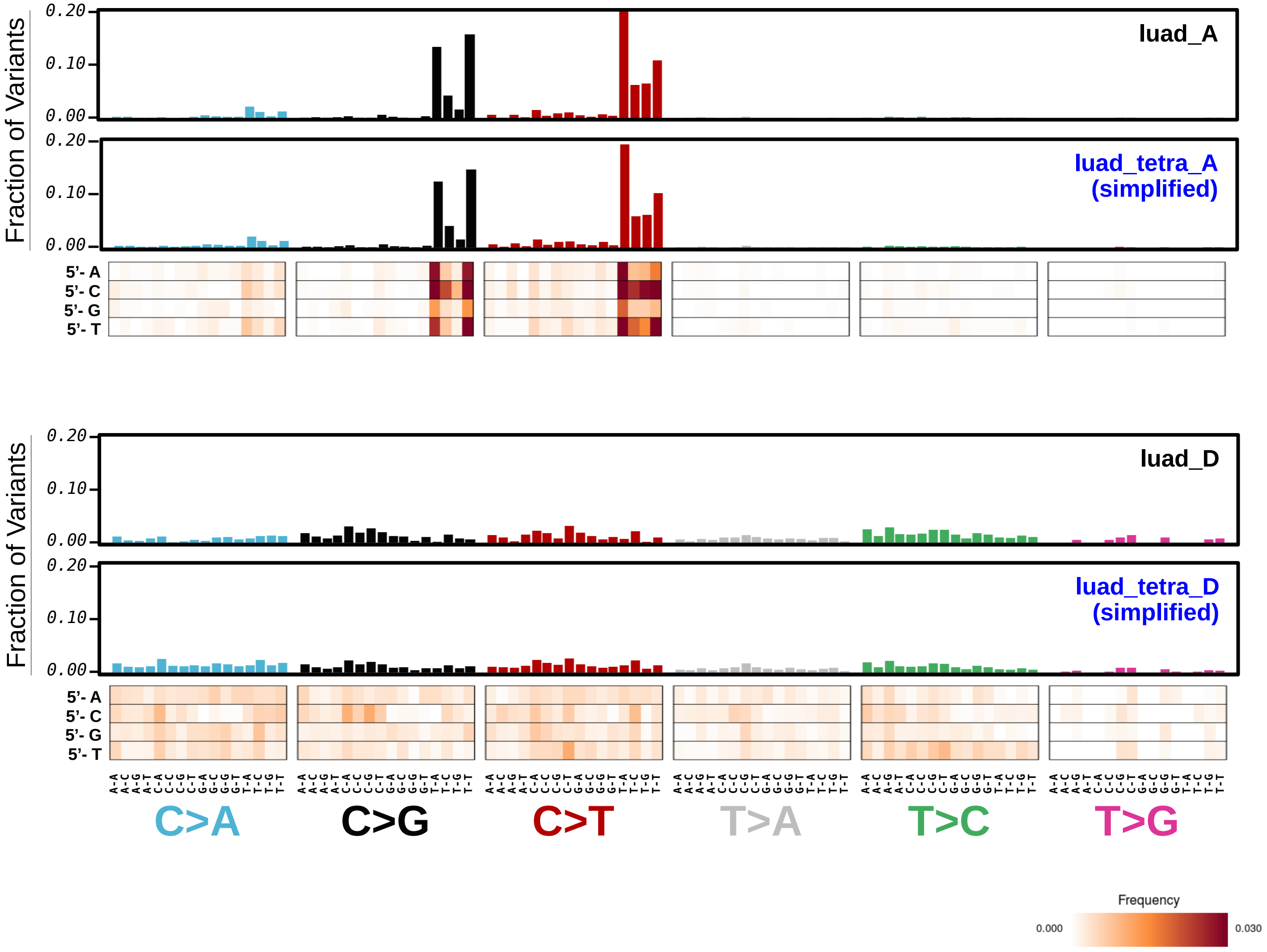


**Supplementary Figure S6.** Mutational signatures extracted from LUAD TCGA. Barplots summarizing the mutational profiles of mutational signatures extracted from TCGA bladder cancer genomes. Tetra-nucleotide mutational signatures were simplified to the corresponding tri-nucleotide mutational pattern by DNA variant aggregation. Corresponding tri- and simplified tetra-nucleotide mutation types were shown as barplots. Heatmaps are visual representations of the tetra-nucleotide mutational signatures, where tri-nucleotide mutation types are shown on the x-axis, and the extra 5'-end nucleotides are shown on the y-axis. Box color intensity tracks with mutation type frequency.

### Supplementary Figure S7

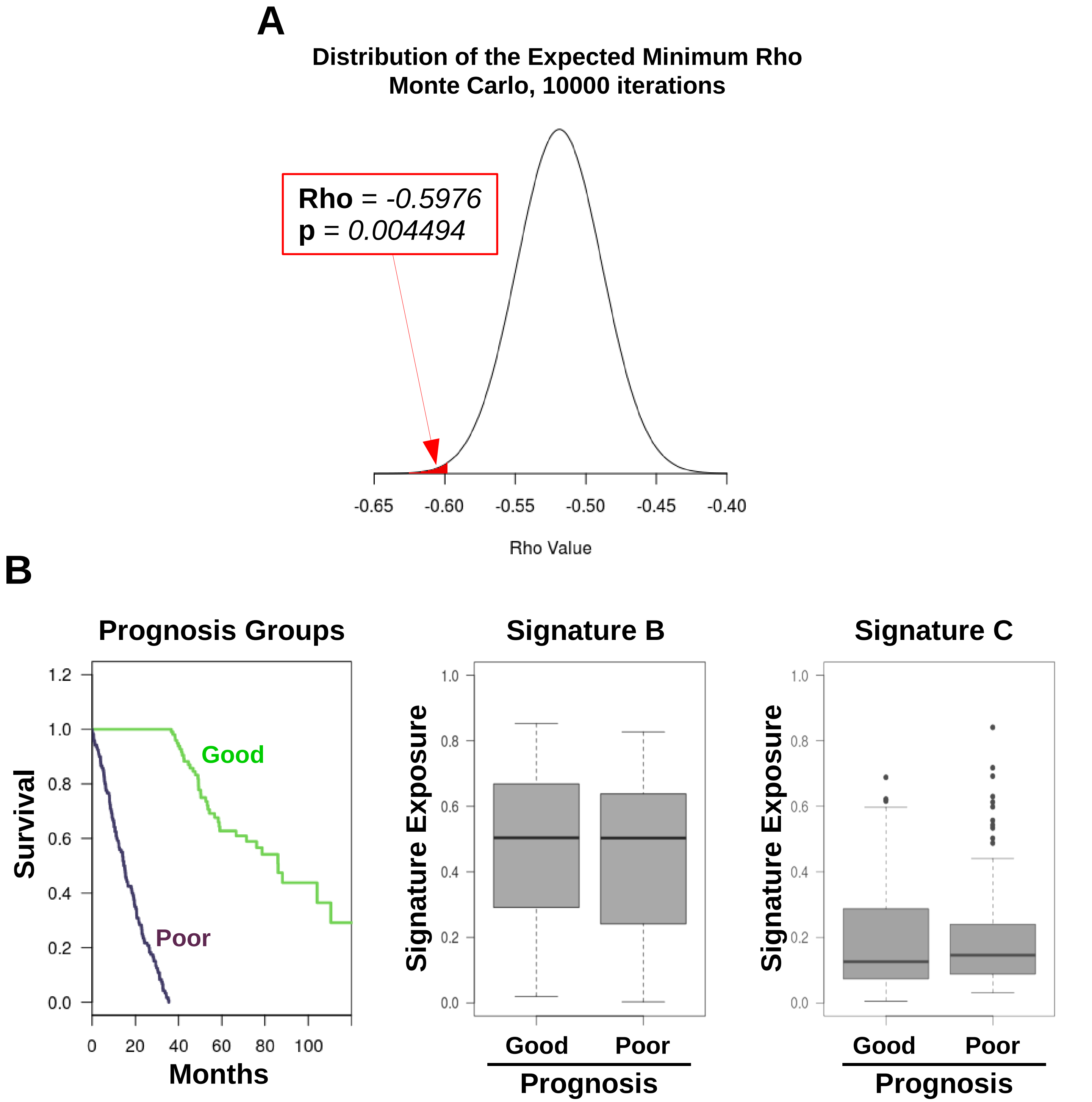


**Supplementary Figure S7.** Correlations among mutational signatures and clinical features. A) Distribution of the minimum correlation coefficients computed by a Monte Carlo simulation (10,000 simulations) to simulate signature exposures. The probability of detecting a minimum coefficient less or equal than the observed Rho for the pair of signatures *luad_B* and *luad_C* (Rho=-0.5976) was p=0.00449 (red portion of the area under the curve). B) Signature exposures were analyzed with respect to prognosis. Survival of the two patient groups (good prognosis vs. poor prognosis) is shown in a survival plot (left). Relative exposures of *luad_B* (middle) and *luad_C* (right) signatures were assessed in both good- and poor-prognosis patients, revealing no differences.
